# Cerebral Encoding of Word Classes is Distributed and Context-Dependent

**DOI:** 10.64898/2026.07.22.739973

**Authors:** Natalia Bekemeier, Marianne Hundt, Zirui Huang, Martina Djordjijevic, Alexis Hervais-Adelman

## Abstract

Word classes such as nouns, verbs, and adjectives are fundamental units of language, but their neural encoding remains unclear. Here, we investigate whether word classes are processed as invariant, context-independent lexical categories or depend on sentence context during language processing. We analyse source-localized magnetoencephalography (MEG) data from 200 native Dutch participants who read or listened to sentences and their scrambled counterparts (word lists) from the MOUS dataset. Time-resolved encoding models are used to predict neural responses from major word classes (noun, verb, adjective) and other linguistic variables including word frequency, surprisal, entropy, word length, and ordinal position. Across modalities, we observe a significant interaction between word class and context (sentences vs. word lists), manifest as dynamic modulation of neural responses in a widespread cortical network including bilateral perisylvian, frontal, and midline regions previously implicated in lexicosemantic, structural, and pragmatic processing. This interaction emerges early and reappears later in processing, with distinct temporal profiles for reading and listening. Within-condition effects reveal that word class contributes to neural responses at the level of individual words, but this contribution is context-dependent: it is robust in sentences across modalities and in word-list reading, but absent in auditory word lists. These results indicate that word-class encoding is shaped by the interaction between word-level properties and sentence context during real-time language processing.

## Introduction

Grammatical categories such as *word classes*, e.g., nouns, verbs, and adjectives, are fundamental to language processing, yet their neural encoding remains poorly understood. Neuroimaging and neuropsychological studies provide partially divergent evidence regarding neural organization of major word classes (nouns, verbs, adjectives). Lesion studies have reported dissociations between noun and verb processing, with damage to the left temporal regions often affecting noun processing and lesions to the left frontal regions impairing verb processing (Miceli et al., 1988; Bates et al., 1991; Damasio and Tranel, 1993; Shapiro and Caramazza, 2003). In contrast, functional neuroimaging studies have yielded heterogeneous results, reporting both overlapping and category-specific activation patterns for nouns and verbs depending on task demands, stimulus types, and languages (Fujimaki et al., 1999; Damasio et al., 2001; Saccuman et al., 2006; Vigliocco et al., 2006; Longe et al., 2007; Momenian et al., 2016). In many cases, observed differences can be attributed to semantic or task-related factors rather than to word class per se (Longe et al., 2007; Saccuman et al., 2006). Together, the reported findings provide evidence for distinctions between such major word classes as nouns and verbs in multiple languages, although the nature of these distinctions remains debated (Vigliocco et al., 2011; Kemmerer, 2014). In contrast, neurobiological data suggesting the existence of adjectives as a distinct word class is relatively limited (Westerlund and Pylkkänen, 2017; Fyshe et al., 2019; Misra et al., 2024).

The apparent discrepancies between lesion-based and neuroimaging results may reflect fundamental methodological differences: lesion studies provide causal evidence about brain regions essential for specific functions, whereas functional imaging yields correlational patterns of co-activation that depend on task demands and experimental design. Furthermore, the ostensible overlap between noun- and verb-related activations reported in previous neuroimaging studies does not preclude fine-grained, category-sensitive subregions within a shared cortical network. Methods with high temporal resolution such as magnetoencephalography (MEG) could capture the rapid dynamics of word-by-word language processing and, thus, help resolve word-class effects. Another possible explanation for the inconsistent findings is that a word class is best conceived of in a coherent context, e.g., in a sentence or a phrase. This problem has received comparatively little attention in neuroimaging studies despite its central role in linguistic theory. Overall, current neurobiological evidence supports a graded and distributed neural organization of word classes with category-specific subregions embedded within a broader, shared semantic and lexical network (Aarts, 2004, 2007; Keizer, 2023).

A central question in linguistic theory is whether word classes reflect intrinsic lexical properties of individual words or emerge from contextual cues during sentence processing. Although words are often associated with a default grammatical category, e.g. *table* - a noun, *run* – a verb, many lexical items can occur with multiple grammatical functions (e.g. *table* as a verb in “to table a motion” or *run* as a noun in “a run on the bank”). In such cases, the word class of a word is shaped by contextual cues and distributional patterns across language use, rather than being regarded as an invariant property of lexical entries (Croft, 2001; Haspelmath, 2007). This variability raises the question whether word class contributes to neural responses independently of context or primarily through its interaction with sentence-level structure. (Dixon, 1982; Schachter and Shopen, 1985; Croft, 2001, 2002; Dixon and Aikhenvald, 2004; Haspelmath, 2007; Bisang, 2010; Shao et al., 2024). Recent developments in philosophy, cognitive science, and linguistics have challenged a strictly categorical view of word classes, instead conceiving of grammatical categories as graded, with flexible boundaries and within-class variability (Aarts, 2004, 2007; Lauwers and Van Goethem, 2020; Keizer, 2023; Shao et al., 2024). Within such a framework, word classes reflect contextual probabilistic tendencies rather than invariant, discrete bundles of features. (Vigliocco et al., 2011; Kemmerer, 2014; Westerlund and Pylkkänen, 2017; Fyshe et al., 2019; Misra et al., 2024).

Despite extensive behavioural and neurobiological evidence, it remains unclear if word class affects neural responses independently of coherent context or primarily through interactions with structured linguistic input. To address this question, the present study investigates the neural encoding of word classes using source-localized magnetoencephalography (MEG), which provides high temporal resolution for tracking word-by-word processing. We analysed data from the Dutch “MOUS” dataset (Schoffelen et al., 2019), containing MEG recordings of 200 participants while they were reading or listening to intact complex sentences and their scrambled counterparts (word lists). This design permits a direct comparison between conditions with and without sentence-level context, allowing us to test whether word-class-related effects generalize across contexts or depend on structural coherence.

We employed encoding modelling (Hoerl and Kennard, 1970; Marquardt and Snee, 1975; Buteneers et al., 2013; Liu and Dobriban, 2019) to predict neural responses from multiple lexical and contextual predictors, including word class (nouns, verbs, and adjectives), lexical frequency, ordinal position, entropy, surprisal, and word length. This approach allows us to determine whether word class explains additional variance in neural activity beyond other theoretical predictors of language processing, following previous work on the same dataset (Huizeling et al., 2022). Specifically, we tested if word-class effects (i) are comparable in sentence and word list conditions, consistent with invariant word-intrinsic lexical encoding; (ii) emerge only in intact sentences, consistent with context-dependent processing; or (iii) differ across conditions, indicating that sentence context shapes cerebral encoding of word class.

## Materials and Methods

### Participants

Two hundred native Dutch speakers (age 18-33 years, mean 22; 100 female) participated in an MEG study of language processing (MOUS dataset (Schoffelen et al., 2019)). Participants received input in either a visual (reading) or auditory (listening) modality. Here, we provide a summary of the methods employed by the authors of the original dataset and the additional steps that were undertaken in the present series of analyses.

### Experimental design and stimuli

Participants read or listened to intact sentences and their scrambled counterparts (word lists), presented in a word-by-word format (reading) or as continuous speech (listening). Each participant was presented with 120 sentences and 120 word lists. Word list stimuli were generated by permuting word order while minimizing local phrase structure building. Attention and comprehension were monitored using intermittent *yes/no* comprehension questions.

### Linguistic predictors

Word class annotations were obtained from precomputed morphosyntactic labels (Huizeling et al., 2022). Analyses were restricted to words tagged as nouns, verbs, and adjectives. Word class was dummy-coded (noun as reference, verb and adjective as binary regressors). Additional predictors included lexical frequency, index (defined as the ordinal position of a word within a sentence), surprisal, entropy, and word length (Supplemental materials A.1). In reading, word length was operationalized as the number of characters, whereas in listening, word length was defined as the duration of a word in milliseconds. Lexical frequency was derived using the NLCOW2012 corpus (Schäfer and Bildhauer, 2012), and a trigram language model was trained on this corpus (Van den Bosch and Berck, 2009) to generate surprisal and entropy values. Lexical frequency transformed to 9+log10(frequency). Surprisal was computed as negative log probability of occurrence of a word in the context of the previous two words. Entropy was defined as the uncertainty over next-word distributions. (Zuur et al., 2010; Cohen et al., 2013; Dormann et al., 2013).

### Data acquisition and analysis

Structural MRI scans were acquired on a 3T Siemens system using a T1-weighted MPRAGE sequence (1 mm isotropic resolution) and used for MEG source reconstruction.

MEG data were recorded with a 275-axial gradiometer (CTF; 1200 Hz sampling rate). Data analyses were run separately for visual and auditory modality, following the analysis pipeline employed by Arana and colleagues (Arana et al., 2020) for pre-processing, source reconstruction and spatiotemporal alignment. Data were band-pass filtered (0.5–20 Hz), epoched relative to word onset, and downsampled to 120 Hz. Artifacts were excluded by marking contaminated samples.

Source reconstruction was performed using linearly constrained minimum variance beamforming (LCMV) (Veen et al., 1997). Source activity was projected onto 382 cortical parcels based on the Conte69 atlas (Van Essen et al., 2012; Schoffelen et al., 2017) (Supplementary materials A.2). Within each parcel, the first five principal components were retained.

### Spatiotemporal alignment

To boost similarities between subject-specific signals in response to the stimuli and to reduce intersubject variability, Multiway Canonical Correlation Analysis (MCCA) (de Cheveigné et al., 2019; Arana et al., 2020) was applied to find projections of the five principal components. MCCA was performed separately for each condition and modality following the implementation of Arana et al. (2020) using their publicly available code (also employed by Huizeling et al. (2022)). We estimated the canonical components out of sample using cross-validation to reduce the risk of overfitting. For each analysis, all trials were partitioned into five folds, estimating MCCA weights on the training folds (96 trials) and subsequently applying them to the test fold (24 trials) to obtain subject-specific canonical components with the same temporal resolution as the original source signals. The resulting canonical time-series were used as the neural response signals for the subsequent time-resolved ridge regression encoding models.

### Encoding models

Encoding models were fitted to canonical components time-locked to word onsets (nouns, verbs, and adjectives). For each participant, parcel, and timepoint, we estimated ridge regression models with five-fold cross-validation (Hoerl and Kennard, 1970). Regularisation parameter was selected from a list λ values (0.002, 0.005, 0.010, 0.020, 0.050, 0.100, 0.200, 0.500, 1.000, 2.000, 5.000). For each λ value, prediction performance was evaluated across all cross-validation folds using mean squared error (MSE), and the λ yielding the lowest average MSE was selected.

To separate the contribution of word class and word class by context interaction from that explained by all other variables, we employed a model comparison scheme previously reported by Huizeling et al. (2022). The model comparison scheme quantified the extent to which a model including a predictor of interest, i.e. the full model, explained variance in the MEG signal, above and beyond a reduced model that did not include the given predictor. Model performance was quantified using coefficient of determination (COD), defined as

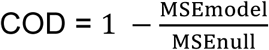

where mean squared error (MSE) between predicted and observed neural responses was computed on held-out data from cross-validation and averaged across folds (Huizeling et al., 2022).

Two complementary model comparison schemes were used to characterize word-class encoding. First, to identify context-dependent modulation of word-class encoding, we compared a full model containing word class by context interaction term against a reduced model containing only the corresponding main effects:

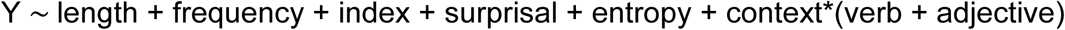

Where context was coded as -1 (word lists) and 1 (sentences) and word class was dummy coded using nouns as the reference category. All predictor variables except binary variables were standardized using z-score. The reduced model excluded the interaction terms:

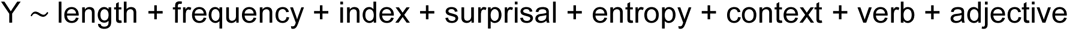

Second, to determine whether word class contributed to the neural responses within each context (sentences vs. word lists), we estimated separate full models for each condition:

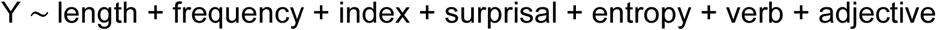

And compared them to reduced models, where word-class dummy variables were dropped:

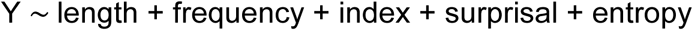

The resulting difference in prediction performance reflects the contribution of word-class predictors within each condition. In both analyses, the contribution of word-class predictors was evaluated above and beyond the variance explained by word length, lexical frequency, ordinal position, surprisal, and entropy. Thus, model comparisons quantified the unique contribution of word class (or its interaction with context) after accounting for these lexical and contextual variables.

For statistical inference, i.e. comparison of the results with the distribution under the null hypothesis, null distributions were obtained by permuting predictor-response mappings within subjects (50 times). The shuffled predictor matrix was refit using the same cross-validation folds and λ-selection procedure as the observed models, both for the word class by context interaction and word class models. The model comparison scheme was also applied to the averaged 50 null distributions. The resulting CODs of the original models and those of the averaged 50 random permutation counterparts (shufCODs) entered statistical analyses.

### Statistical analysis

Statistical inference was performed across subjects using non-parametric permutation-based dependent-samples t-tests. We compared the observed CODs to their null distribution counterparts (shufCODs). Multiple comparisons across time were controlled using a max-statistic approach (5,000 permutations). Spatial correction across parcels was performed using Benjamini-Hochberg false discovery rate (FDR) correction.

### Post-hoc analyses

Post-hoc analyses were performed only within parcels showing a significant word-class effect within conditions (sentences vs. word lists). To assess word-class-specific effects, regression coefficients for word class predictors were compared using non-parametric paired t-tests (verb vs. noun, adjective vs. noun, verb vs adjective) at each timepoint. Multiple comparisons across time were corrected using the max-statistic method, and pairwise contrasts were adjusted with Bonferroni correction (α = 0.05/3 ≈ 0.0167). Spatial correction across parcels was performed applying FDR correction.

## Results

We first tested whether the contribution of word class depended on sentence context (Figure 1), then examined whether word-class predictors explained additional neural variance within each condition (Figure 2), and finally identified which word classes contributed to these effects (Figures 3–6).

**Figure 1.**
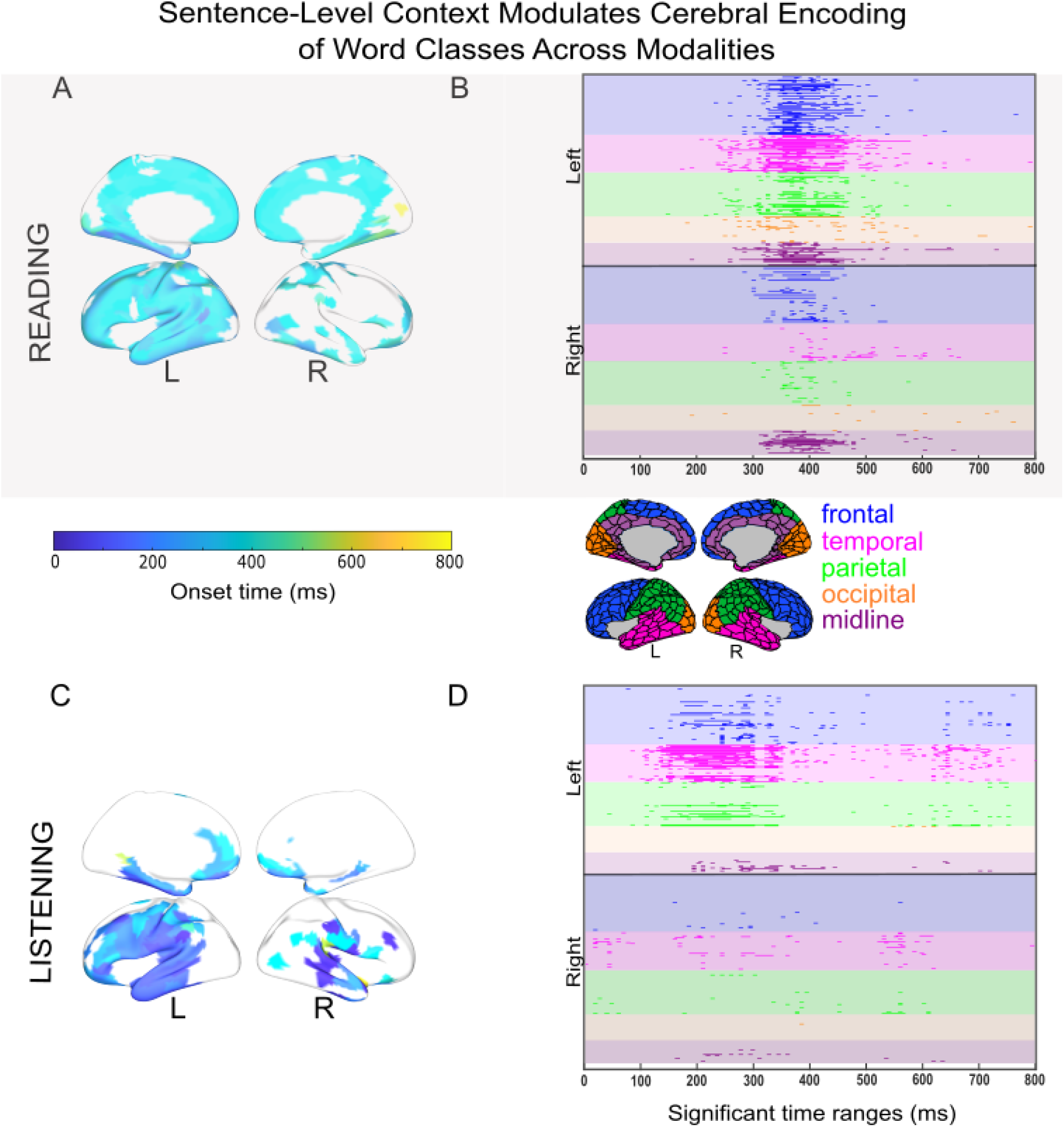
Sentence-level context modulates cerebral encoding of word classes across modalities. Significant word class by context interactions were identified by comparing the full encoding model with a reduced model that excluded the word class by context interaction term, while controlling for word class, word length, lexical frequency, ordinal position, surprisal, and entropy (temporal correction using the max-statistic approach and spatial correction using FDR (q < 0.05)). Results for reading are displayed in the upper panels (A, B; light-gray background) and for listening in the lower panels (C, D). Panels (A, C) show cortical maps indicating the earliest significant time point of the effect (0-800 ms post word-onset). Raster plots in panel (B, D) show the temporal and spatial extent of effects across parcels. Each row represents an atlas parcel, and dark bands indicate time windows where the effect reached significance. Parcels (total: 382; 191 per hemisphere) are grouped by hemisphere and neuroanatomical subdivision, colours as shown on Atlas. The interaction emerged early in both modalities and reappeared at later latencies, indicating that sentence-level context dynamically shapes neural responses to word classes.

**Figure 2.**
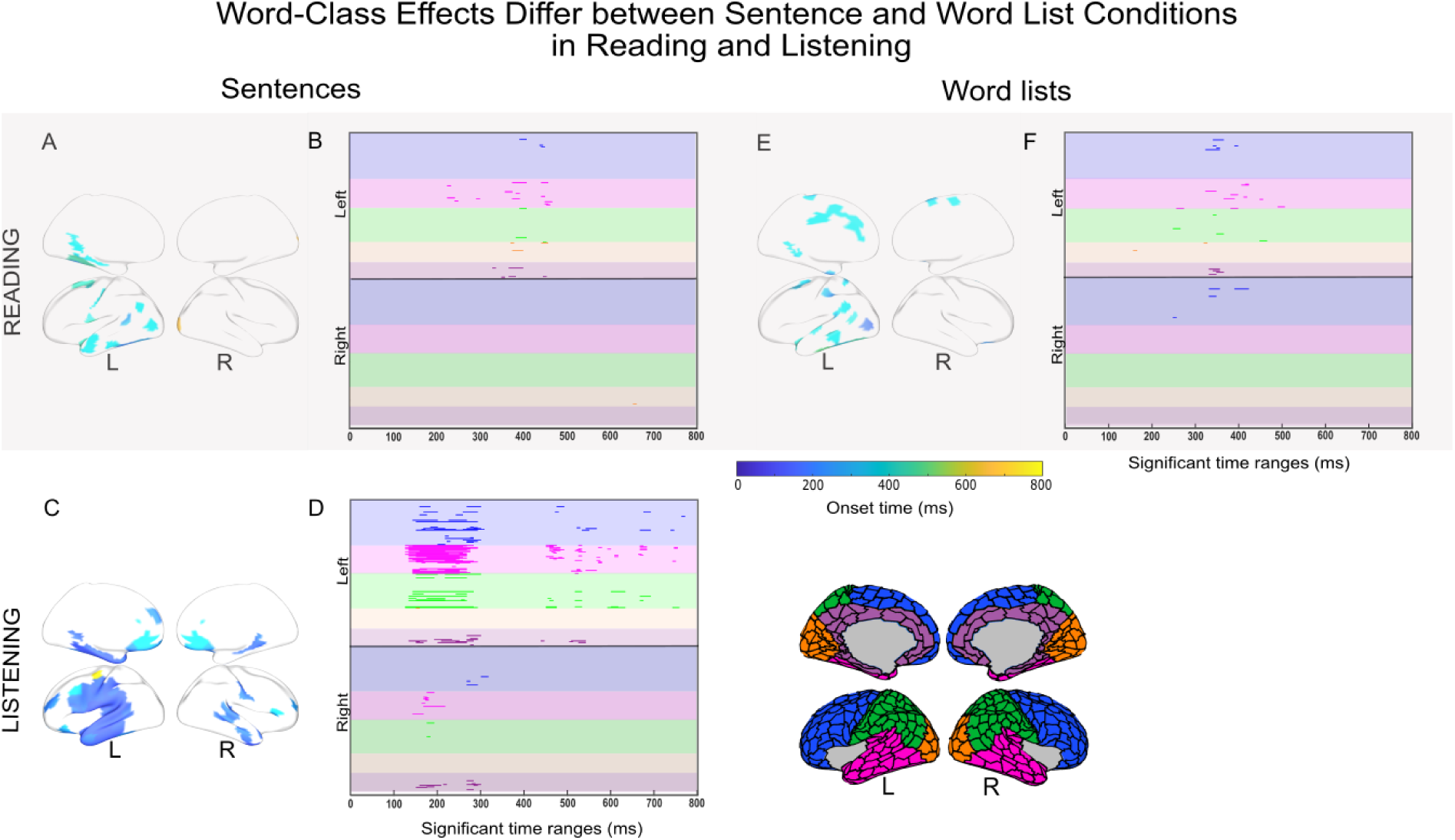
Neural encoding of word class differs between sentence and word-list conditions in reading and listening. Significant word-class effects were identified by comparing the full encoding model with a reduced model lacking the word-class predictors, while controlling for other lexical parameters (temporal correction using the max-statistic approach and spatial correction using FDR (q < 0.05)). Word-class effects in sentence conditions are shown in the left panels; the effect of word class in word-list reading is shown in the right panels. No significant word-class effects were observed in word-list listening. Results for reading are displayed in the upper panels and for listening in the lower panels. Panels (A, C, E) show cortical maps of the earliest significant time point of the effect (0-800 ms; dark blue to yellow). Raster plots in panel (B, D, F) show the temporal and spatial extent of effects across parcels. Each row represents an atlas parcel and dark bands indicate time windows where the effect reached significance after. Parcels (total: 382; 191 per hemisphere) are grouped by hemisphere and neuroanatomical subdivision, colours as shown on Atlas.

### Interactions across modalities

After accounting for word length, lexical frequency, ordinal position, surprisal, and entropy, the inclusion of the word class x context terms into the encoding model explained additional variance in neural responses across a widespread bilateral cortical network. These effects indicate that the neural encoding of word class depends on whether words are embedded in a coherent sentence context or presented as isolated sequences. The interaction effects emerged early, with onsets around 200 ms in reading (Figure 1A, B) and immediately following word onset in listening (Figure 1C, D), initially involving temporal and parietal areas before extending to the frontal and midline cortices (Figure 1B, D). The temporal dynamics differed between modalities, with peak interaction effects occurring around 400 ms in reading and 250 ms in listening. Although most pronounced between 200-400 ms after word onset, interaction effects re-emerged at later latencies (>500 ms), indicating that contextual modulation of word-class encoding is dynamically updated throughout language processing rather than restricted to a single processing stage. In listening, the later interaction comprised temporally distinct right-hemispheric (peak: ∼550 ms) and left-hemispheric (peak: ∼650 ms) effects. Together, these findings show that word class contributes to neural responses but that its expression is dynamically shaped by context across modalities.

### Within-condition effects

To determine whether word class explained additional variance in neural responses within each condition, beyond the contribution of word length, lexical frequency, ordinal position, surprisal, and entropy, we examined word-class effects separately within sentence and word list conditions in each modality. In reading, sentences (Figure 2A, B) and word lists (Figure 2E, F) elicited significant word-class effects in a constrained but comparable number of brain areas. The significant areas were predominantly left-lateralized and largely non-overlapping, suggesting partially distinct neural mechanisms associated with word-class encoding in reading in the presence and absence of sentence context.

In listening, sentence processing elicited more widespread word-class effects than sentence reading (Figure 2C, D). These effects emerged early and were most prominent between 125-300 ms after word onset, involving predominantly left-hemispheric regions but also several right hemisphere areas, including the right superior temporal gyrus, pars triangularis, orbitofrontal area, and anterior cingulate cortex. Word-class effects restricted to the left hemisphere emerged later (400-800 ms) in temporal regions and extended to frontal, parietal, midline structures, and medial temporal lobe regions. The involvement of these regions likely reflects the incorporation of word class into higher-order semantic and syntactic representations during ongoing sentence interpretation. In contrast, no reliable contribution of word-class predictors was observed during listening to word lists.

Together, the within-condition results show that word class significantly predicts the word-by-word elicited neural responses during language processing but that the magnitude and the distribution of this effect strongly depends not only on the presence of sentence-level (or phrasal) context but also on modality.

### Post-hoc pairwise contrasts: Category-specific effects

To determine which word classes drove the word-class effects identified in the previous analysis, we conducted targeted pairwise contrasts on regression coefficients. The model used nouns as the reference level; accordingly, coefficients for verbs and adjectives reflect their respective differences relative to nouns, while the verb-adjective contrast was derived by comparing the corresponding coefficients (Figures 3-6). These contrasts indicate time- and parcel-specific differences in neural responses associated with individual word classes after accounting for word length, lexical frequency, ordinal position, surprisal, and entropy. For visualization and reporting, effects are described as noun-, verb-, or adjective-dominant when a given category significantly differed from at least one other category. Parcels showing significant effects across multiple pairwise comparisons were classified as exhibiting sensitivity to multiple word-class contrasts.

In sentence reading, three brain areas showed sensitivity to specific word-class contrasts (Figure 3). Parcels of the left middle temporal and inferior temporal gyri (Figure 3A, D) were selectively sensitive to adjective-dominant contrasts, consistent with the involvement of ventral temporal cortex in fine-grained semantic feature processing. The parcel in the left middle temporal gyrus was sensitive to adjective over verb around 400 ms post word-onset (Figure 3A). In the later time window (>770 ms) the parcel in the left inferior temporal gyrus showed sensitivity to adjective over verb. In contrast, a parcel of the left temporal pole showed selective sensitivity to verb-dominant contrasts (Figure 4C) in the time window of 400-450 ms.

**Figure 3.**
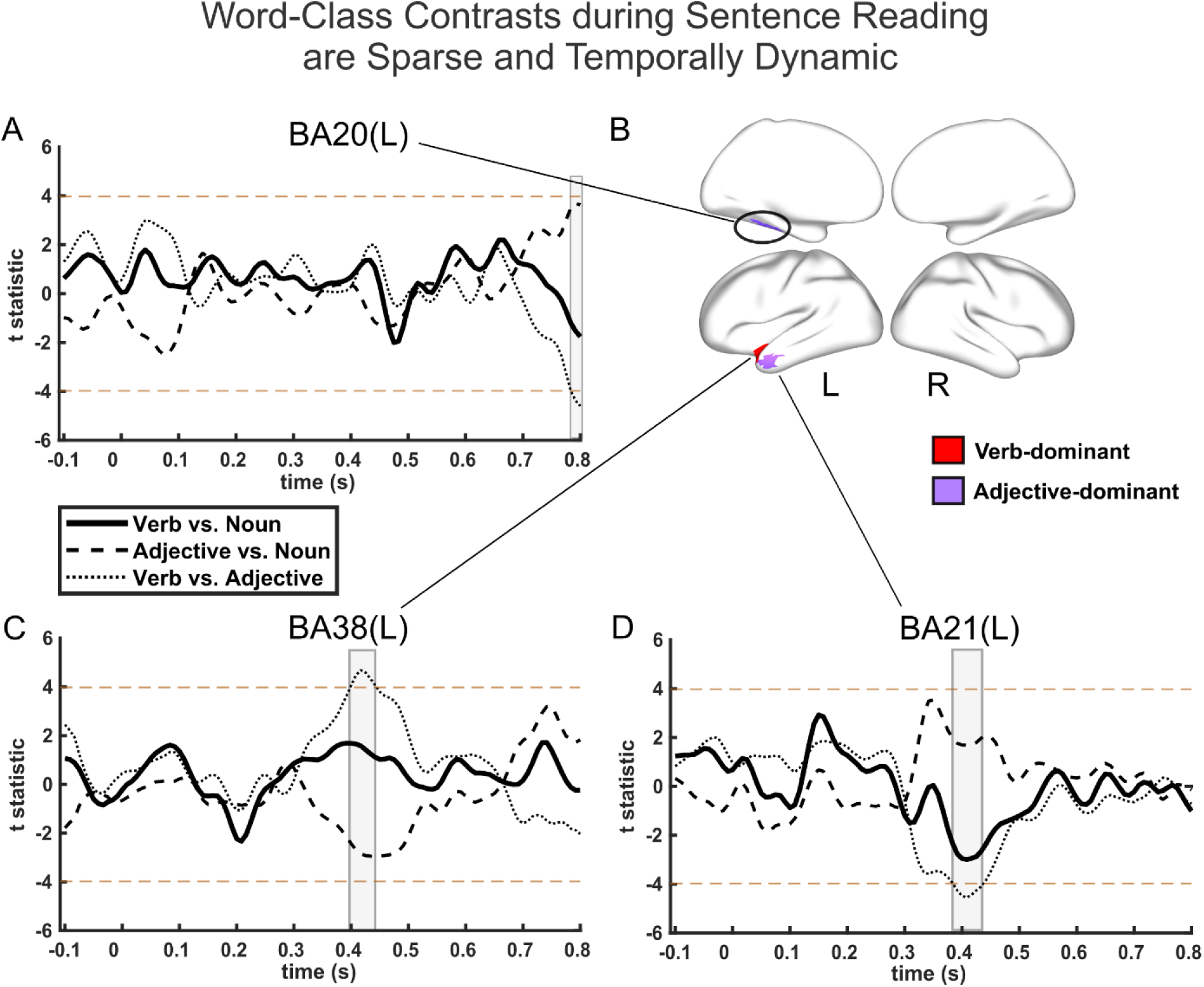
Sentence reading reveals spatially restricted and temporally dynamic sensitivity to word-class contrasts. Pairwise comparisons of regression coefficients identified which word-class contrasts contributed to the significant word-class effect shown in Figure 2. Panels (A, C, D) display time courses of t-statistics for pairwise word-class contrasts: verb vs. noun (solid), adjective vs. noun (dashed), and verb vs. adjective (dotted). Horizontal dashed brown lines indicate the significance threshold. Transparent bars indicate time windows where the effect reached significance after temporal correction using the max-statistic approach and spatial correction using FDR (q < 0.05). Panel (B) shows cortical parcels exhibiting significant word class sensitivity in the sentence condition. Parcels are classified as verb-dominant (red) and adjective-dominant (purple). The example waveforms illustrate adjective-dominant contrasts in the left middle temporal gyrus (panel A, called out from the upper left cortical map in panel B) and in the left inferior temporal gyrus (panel D, called out from the bottom left cortical map in panel B), and a verb-dominant contrast in the left temporal pole (panel C, called out from the bottom left cortical map in panel B).

In word-list reading, category-specific effects were scarce and more heterogeneous. One parcel of the left primary somatosensory cortex (Figure 4C) exhibited sensitivity to noun over verb at 366-410 ms. Two additional parcels located in the left auditory cortex and left ventral anterior cingulate cortex showed dynamic sensitivity to multiple word-class contrasts. The parcel in left ventral anterior cingulate cortex (Figure 4A) was sensitive to verb over noun in the pre-stimulus onset window (-40 – -16 ms) followed by the sensitivity to noun over verb (310-433 ms) and adjective over verb (316-390 ms). The parcel in the left auditory cortex (Figure 4D) was sensitive to verb over noun in a very early time window (-8 – 20 ms), followed by a later sensitivity to noun over verb at 240-275 ms. Crucially, many parcels in both sentence and word list conditions did not exhibit selectivity for any word-class contrasts. These findings indicate that the neural encoding of word class during reading is often multiplex and dynamic, rather than statically tied to a single word class.

**Figure 4.**
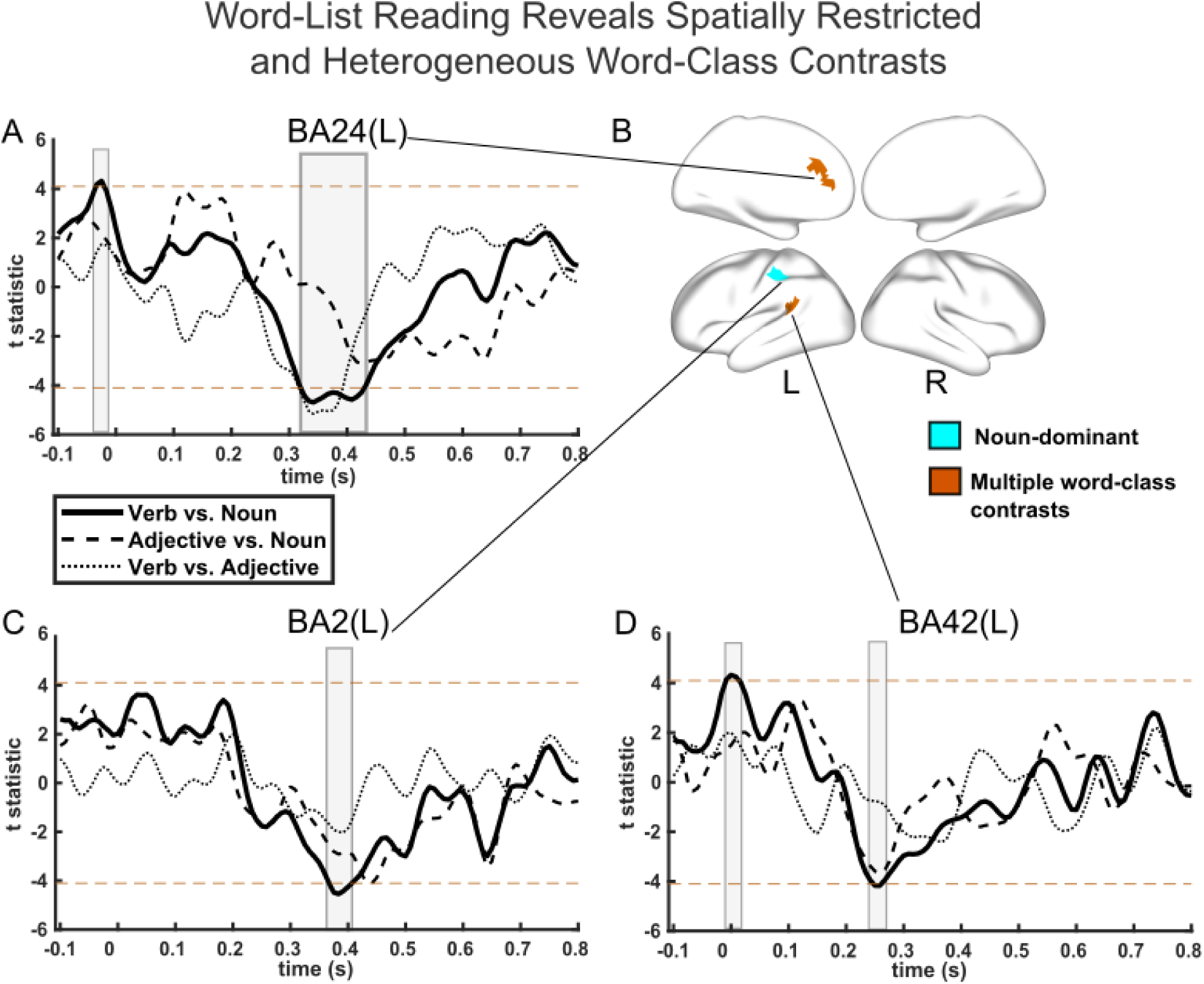
Word-list reading elicited sparse and heterogeneous sensitivity to word-class contrasts. Pairwise comparisons reveal which word classes contributed to the word-class effect during word-list reading. Panels (A, C, D) show time courses of t-statistics for pairwise word-class contrasts: verb vs. noun (solid), adjective vs. noun (dashed), and verb vs. adjective (dotted). Horizontal dashed brown lines indicate the significance threshold. Transparent bars indicate time windows where the effect reached significance after temporal correction using the max-statistic approach and spatial correction using FDR (q < 0.05). Panel (B) shows cortical parcels exhibiting significant word-class contrasts in the word list condition. Parcels are classified as noun-dominant (cyan) or exhibiting dynamic sensitivity to multiple word-class contrasts (brown). Most significant parcels showed temporally varying differences between multiple word-class contrasts rather than a persistent preference for a single word-class contrast (panels A and D). Noun-dominant sensitivity is demonstrated in the left posterior primary somatosensory cortex (panel C).

Unlike reading, where word-class-specific effects were sparse, sentence listening revealed a broader distribution of pairwise word-class contrasts. Category-specific sensitivity was observed across a bilateral network of cortical regions, with spatially distinct patterns of sensitivity to different pairwise word-class contrasts (Figure 5). Category-specific effects occurred across early and late processing windows, with most effects clustering between 100-350 ms and from approximately 450 ms until the end of the epoch (Figure 6).

**Figure 5.**
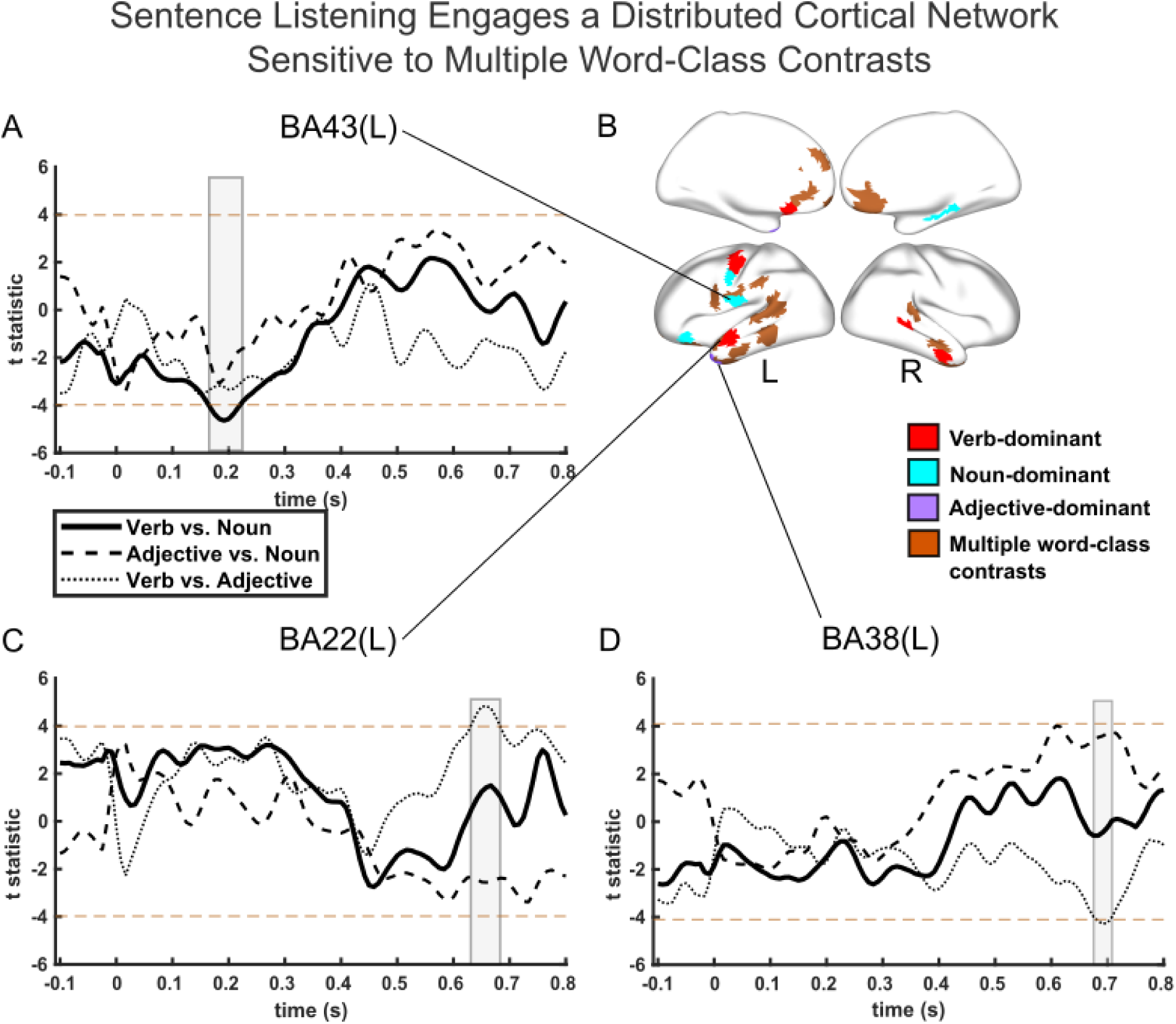
Sentence listening reveals a distributed cortical network sensitive to multiple word-class contrasts. Sentence listening engaged a broader and more distributed network than reading, with both category-specific sensitivity and sensitivity to multiple word-class contrasts. Panels (A, C, D) show representative parcel-level time courses of t-statistics for pairwise word-class comparisons during sentence listening: verb vs. noun (solid), adjective vs. noun (dashed), and verb vs. adjective (dotted). Horizontal dashed brown lines indicate the significance threshold. Transparent bars indicate time windows where the effect reached significance after temporal correction using the max-statistic approach and spatial correction using FDR (q < 0.05). Panel (B) displays cortical maps summarizing parcels with significant word-class-related sensitivity, classified as verb-dominant (red), noun-dominant (cyan), adjective-dominant (purple), or showing dynamic sensitivity to multiple word-class contrasts (brown). Example parcels include the left gustatory cortex (A) for a noun-dominant contrast, the left superior temporal gyrus (C) for the verb-dominant contrast, and the left temporal pole (D) for the adjective-dominant contrast.

Noun-dominant contrasts elicited selective sensitivity in four parcels (Figure 5A, B; Figure 6C): noun over adjective in the left orbitofrontal area (8-35 ms), and noun over verb in the left primary gustatory cortex (158-233 ms: Figure 5A), left primary motor cortex (300-360 ms), and the right ventral entorhinal cortex (433-575 ms). Verb-dominant contrasts were observed in six parcels (Figure 5B, C; Figure 6A), including the left primary motor cortex (two parcels), superior temporal gyrus, ventromedial prefrontal cortex, and the right superior and middle temporal gyri. A parcel in the left temporal pole (Figure 5B, D; Figure 6B) exhibited selective sensitivity to adjective over verb at 670-710 ms.

**Figure 6.**
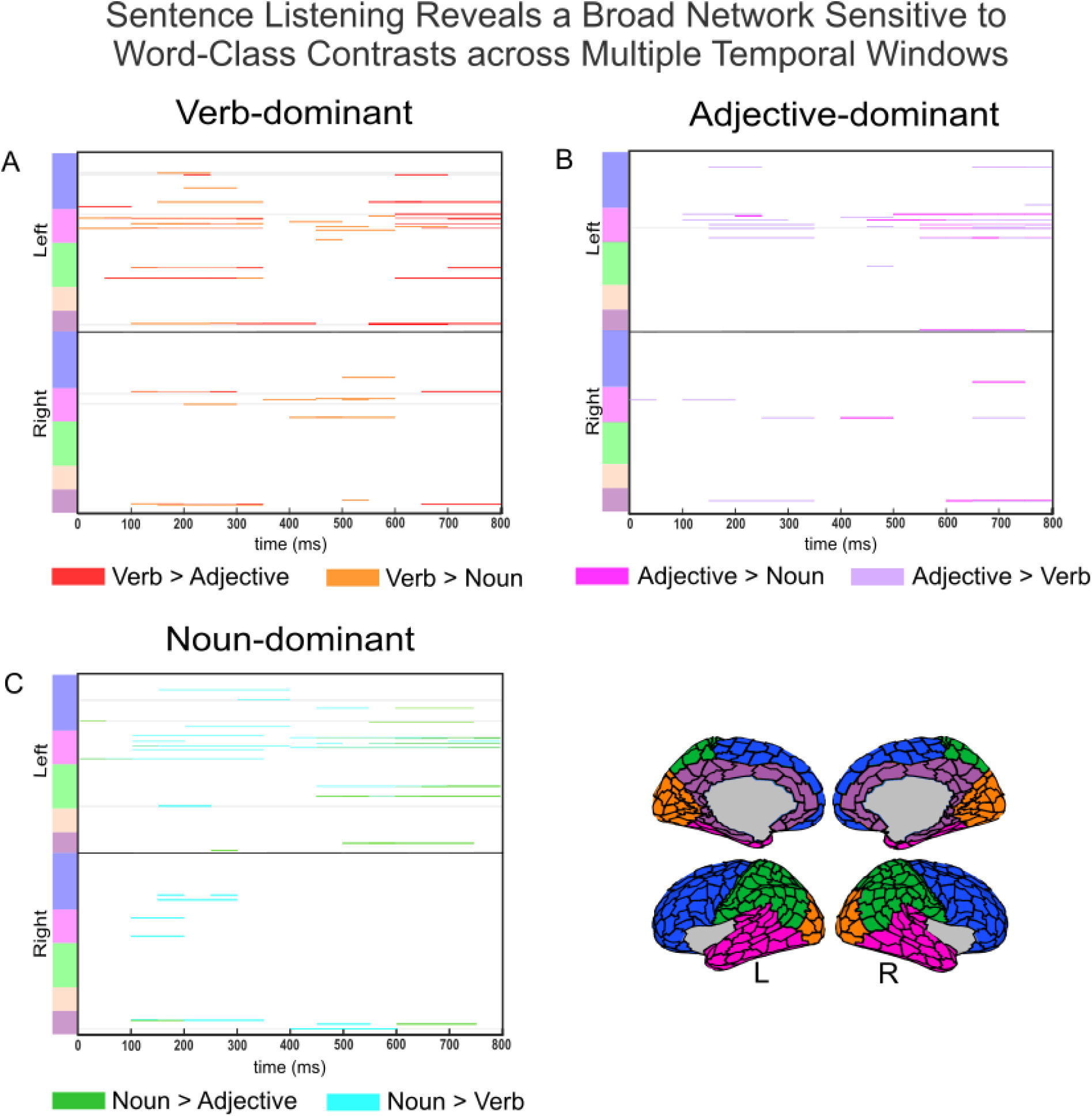
Temporal and spatial distribution of word-class-specific sensitivity during sentence listening. Raster plots in panels (A-C) show the temporal and spatial extent of significant pairwise word-class contrasts across all word-class-sensitive parcels. Each row represents an atlas parcel. Parcels (total: 382; 191 per hemisphere) are grouped by hemisphere and neuroanatomical subdivision, colours as shown on Atlas. Coloured bands indicate time intervals during which specific word-class contrasts reached significance after temporal correction using the max-statistic approach and spatial correction using FDR (q < 0.05): verb > adjective (red), verb > noun (orange), adjective > noun (magenta), adjective > verb (purple), noun > adjective (green), and noun > verb (cyan). Light-gray bands and triangles mark the parcels with word-class-specific sensitivity. The raster plots demonstrate two principal periods of word-class discrimination (approximately 100–350 ms and >450 ms), with extensive temporal overlap among different pairwise contrasts.

In addition to selective responses for individual pairwise word-class contrasts, 23 parcels demonstrated sensitivity to multiple pairwise contrasts (Figure 5B), indicating distributed and temporally dynamic encoding of word class during sentence processing.

### Modality-independent effects

To determine whether word-class-by-context effects generalized across modalities, we identified cortical regions showing overlapping significant interaction effects between reading and listening. Significant overlap was observed across a broad bilateral network, including regions along the superior and middle temporal gyri and ventral frontal areas (Figure 7). This cross-modal convergence indicates that context-dependent modulation of word-class encoding engages partially shared cortical systems during language processing across input modalities. Since modality-independent word-class effects within individual conditions were only observed for sentence conditions and largely overlapped with the word class x context interaction, we focused the cross-modal analysis on the interaction effect only.

**Figure 7.**
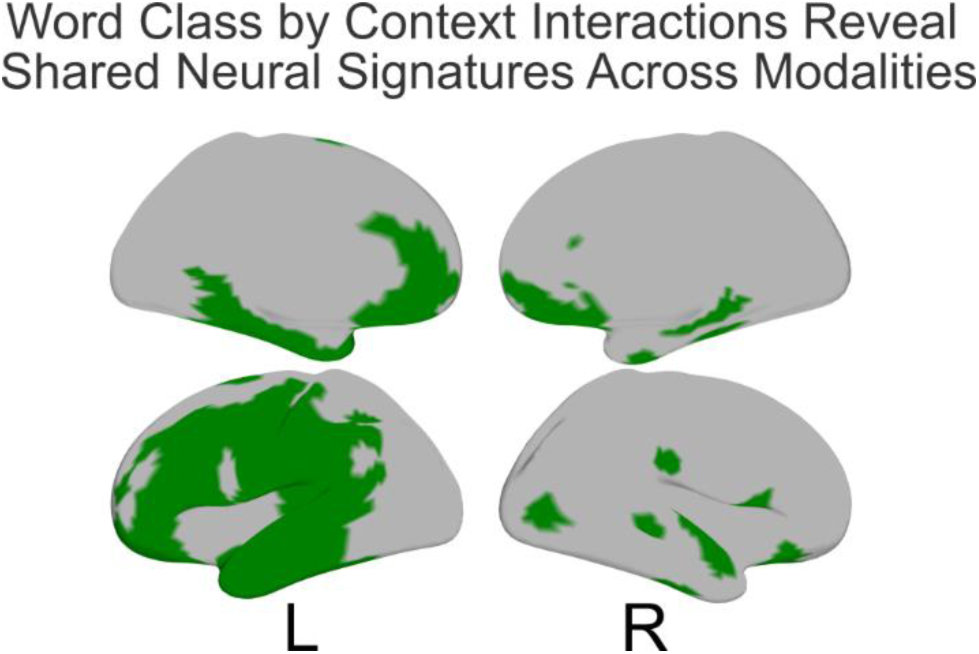
Modality-independent cortical network supporting contextual modulation of word class. Conjunction analysis showing parcels exhibiting a significant word class by context interaction in both reading and listening (after temporal correction using the max-statistic approach and spatial correction using FDR (q < 0.05)). Shared effects were concentrated in bilateral temporal and frontal regions, indicating modality-independent contextual modulation of word-class representations.

## Discussion

A long-standing question in linguistics and cognitive neuroscience is whether word class is represented as a stable lexical property or emerges in sentence context. The present study addressed this question by examining source-localized MEG in native Dutch speakers. Using encoding models, we tested whether word class explains additional variance in word-by-word elicited neural responses beyond lexical frequency, surprisal, entropy, ordinal position, and word length, and whether this contribution depends on coherent sentence context. Across modalities, we observed a spatially and temporally extensive interaction between word class and context, with particularly strong effects in reading. The early onset and later reappearance of this interaction indicate that contextual modulation of word-class encoding unfolds dynamically over time rather that reflecting a single processing stage. Within-condition analyses further showed that word class explained additional variance in neural responses to individual words, with robust word-class effects in sentence processing across both modalities and in word-list reading, but not in word-list listening. The interaction between word class and context was also the most consistent effect across modalities, engaging a widespread bilateral cortical network, including bilateral perisylvian, frontal, and midline regions, that substantially exceeded the spatial extent of within-condition word-class effects. Together, these findings suggest that word class is not expressed primarily as a fixed lexical feature that is retrieved during word recognition. Instead, it appears to be dynamically shaped during incremental processing through interactions between lexical representations and sentence context.

Early recruitment of the temporal and parietal regions likely reflects ventral-stream contributions associated with rapid lexical access and the activation of the word class as probabilistic constraints on word identity, e.g., in the left middle temporal areas (Indefrey and Levelt, 2004; Bemis and Pylkkänen, 2011; Rice et al., 2015; Brennan and Pylkkänen, 2017). At this stage, word-class effects appear closely tied to word-level representations and can be observed not only in the sentence condition across modalities but also in the word-list reading. In contrast, later effects, involving frontal and midline regions, are more consistent with dorsal-stream engagement supporting context-dependent reinterpretation and structural integration (Hickok and Poeppel, 2007; Price, 2010; Saur et al., 2010; Price, 2012; Kemmerer, 2019). Since the dominant effect in the present study is the interaction between word class and context, these later effects may reflect increasing top-down contextual modulation of word-class representations. Thus, the neural encoding of word classes appears to be repeatedly updated as language processing unfolds, transitioning from a ventrally supported lexical access to a dorsally mediated structural interpretation. This dynamic modulation provides a more precise interpretation of dual-stream models of language processing (Hickok and Poeppel, 2000; Friederici, 2002; Hickok and Poeppel, 2004; Friederici, 2012), although the present data do not directly establish a processing hierarchy. Rather than reflecting a strict division of labour, these dynamics suggest that ventral and dorsal systems contribute at different timescales during language processing, in line with hierarchical and predictive accounts of language processing, wherein linguistic information is progressively transformed across interacting representational levels (Friederici, 2012; Heilbron et al., 2022).

Recent studies (Goldstein et al., 2024; Goldstein et al., 2025) propose a hierarchical organization of speech and language processing in which cortical systems largely correspond to distinct levels of a unified acoustic-to-speech-to-language embedding space. In this framework, speech-level embeddings better capture cortical activity the left precentral gyrus and superior temporal cortex, whereas inferior frontal, posterior temporal, and parietal regions are more strongly aligned with higher-level language embeddings. The widespread interaction observed in the present study supports the view that word-class-related effects do not map onto a single representational level within this hierarchy but instead reflect a dynamic shift in the relative contribution of these levels over time and across contexts, consistent with a transition from lexical access to structure-dependent integration during language processing.

Modality-specific differences further contextualize these findings. Listening to sentences elicits earlier and more temporally extended word-class effect than sentence reading, consistent with the continuous and temporally structured nature of the auditory input (Friederici, 2002, 2012). In contrast, word-by-word reading yields later and more spatially restricted word-class effects in both sentence and word list conditions. Crucially, despite these modality-specific differences in timing and spatial distribution, the critical interaction between word class and context is present in both modalities (Figure 7). This convergence suggests that the input modality does not fundamentally alter the nature of word-class representations but rather modulates the temporal dynamics through which lexical information is integrated into and is modified by the unfolding context. Across reading and listening word class is continuously shaped by sentence structure, reinforcing the view that word class is not an invariant lexical property but is dynamically resolved during language processing.

Post-hoc analyses reveal heterogeneous sensitivity to word-class contrasts across cortical regions and time. While some parcels show preferential sensitivity to specific word-class contrasts (e.g., verb-dominant, adjective-dominant effects), most word-class-sensitive areas exhibit dynamic sensitivity to multiple categories. This pattern suggests that category-specific effects are not spatially fixed but instead unfold dynamically during processing. Prior work has reported category-sensitive cortical responses during semantic composition in the left temporal and inferior frontal regions (Bemis and Pylkkänen, 2011; Brennan and Pylkkänen, 2017; Zhang and Pylkkänen, 2018; Matchin et al., 2019), supporting the view that word-class-related information is distributed across cortical networks. The present results extend these findings by showing that such sensitivity is not stable across time but is dynamically constructed during language processing. A notable temporal feature in the post-hoc analyses is the early emergence of verb-dominant contrasts, particularly in the auditory modality. This pattern is broadly consistent with previous findings showing early verb-related effects in rapid sentence processing paradigms, including the structure superiority effect observed using rapid parallel visual presentation (RPVP) (Fallon and Pylkkänen, 2024). Although the paradigms differ substantially in input structure and task demands, both suggest that verb-related information can exert early influence on sentence-level processing. However, these differences preclude strong inferences about shared mechanisms across studies. A possible interpretation of the early verb-dominant sensitivity in sentence listening could be the pivotal role of verbs in constructing sentence representations, as proposed in verb-valency accounts, where verbs constrain or guide the assignment of argument structure.

From a theoretical linguistics perspective, the present findings contribute to ongoing debates about whether word classes are best conceived of as inherent lexical properties or emerge from the interaction between lexical items and the context that they appear in. Traditional accounts assume a relatively stable category membership, whereas distributional and constructionist approaches argue that word classes arise from patterns of use and the interaction between lexical items and grammatical structures (Croft, 2001, 2022). The robust word class by context interaction observed across modalities suggests sensitivity of word-class effects to ongoing integration processes. This does not directly distinguish between representational accounts of lexical categories but rather indicates that the neural correlates of word classes are not fixed across processing stages or contexts. Importantly, this raises the question of whether word classes should be treated as discrete and stable units at the level of processing. While word class was operationalized categorically in the present study, the observed temporal variability suggests that the stability of the category itself may depend on the processing stage and contextual constraints. In this sense, the present results align with construction grammar approaches (Croft, 2001; Croft, 2008; Croft, 2022), in which a word class emerges from the interaction between lexical and grammatical information, rather than reflecting static, context-invariant features.

Several limitations should be noted. The study was conducted in Dutch, which is a verb-second (V2) language; therefore, the observed temporal dynamics may partly reflect language-specific word order properties and may not generalize to languages with different syntactic structures. The word list auditory stimuli differed from sentences not only in syntactic structure but also in prosody and temporal coherence, possibly affecting the processing at multiple levels (Schoffelen et al., 2019; Slaats et al., 2023).The word-by-word presentation constrained naturalistic reading, while preserving incremental processing, which is known to support predictive interpretation of the upcoming input (Wang et al., 2023). The block design may have encouraged task-specific strategies, although it is unlikely that such strategies would differentially affect brain responses to specific word classes. Because the study relied on an existing dataset, experimental manipulations of syntactic complexity or semantic ambiguity were not possible. Finally, source-localized MEG was summarized using a cortical parcellation scheme based on an anatomical atlas, improving interpretability but reducing spatial granularity. Conservative correction for multiple comparisons in parcel-level analyses may have obscured fine-grained effects.

Importantly, the present findings do not establish word-class effects in isolation, as such effects have previously been reported (Vigliocco et al., 2011; Kemmerer, 2014). Instead, they specify the conditions under which word-class effects are expressed in neural responses. Specifically, word class contributes to word-level processing while being systematically modulated by sentence structure and input modality. The robust word class by context interaction identifies a distributed cortical network involved in integrating word class with sentence-level context across reading and listening. This perspective has implications for computational models of language processing, but more broadly for theoretical accounts of word classes. An important aspect of the present approach is that word-class effects were assessed after accounting for multiple correlated lexical predictors, including lexical frequency, surprisal, entropy, word length, and ordinal position. Consequently, the reported effects cannot be explained solely by these established predictors of neural responses to words but instead reflect the unique contribution of word class and its interaction with sentence context.

In conclusion, word class in the human brain is expressed at the level of individual words but is strongly shaped by contextual and modality-dependent processing dynamics. Rather than reflecting either purely word-intrinsic or purely context-dependent properties, word-class effects arise from the interaction between lexical category information and structured linguistic input. Future work employing more naturalistic paradigms, cross-linguistic comparisons, and multimodal imaging will be crucial for further investigations of how word classes emerge in real time during language processing, including both stimulus-locked responses and the ongoing brain states associated with contextual integration during sentence construction.

## Supporting information

Supplemental Materials Summary

Parcel Labels Ordered by Lobe and HM

Parcel Labels

Brodmann Area by Parcel Index

Figure5 Sourcedata

Figure4 Sourcedata

Figure3 Sourcedata

Figure2 Sourcedata for Sentence Reading

Figure2 Sourcedata for Sentence Listening

Figure2 Sourcedata for Wordlist Reading

Figure1 Sourcedata Interaction in Listening

Figure1 Sourcedata Interaction in Reading

## Author Contributions

N.B.: data curation, analysis, investigation, visualization, methodology, writing – original draft, writing – review and editing; A.H.-A.: conceptualization, investigation, methodology, writing – review and editing; M.H.: conceptualization, writing – review and editing; Z.H., and M.D.: writing – review and editing.

## Data Availability

The electrophysiological data analysed in the present study are part of the MOUS dataset (Schoffelen et al., 2019) that has been deposited at https://doi.org/10.34973/37n0-yc51 and is publicly available.

The MEG preprocessing, LCMV beamforming, and multiset canonical correlation analysis (MCCA) follow the analysis pipeline published by Arana et al. (2020; https://doi.org/10.34973/tf8r-rq72).

The subsequent analysis steps are implemented in our own scripts that are openly accessible at https://github.com/bekemeier/Paper_PoS.

## Acknowledgments

This work was supported by the Swiss National Science Foundation (Grant No. CRSII5_209413 awarded to M.N and A.H.-A.). Lexical parameter metrics of the stimuli were kindly shared with us by Eleanor Huizeling and colleagues (Huizeling et al. (2021)). We thank Jan-Mathijs Schoffelen for assistance in adapting portions of the published analysis code from Arana et al. 2020.

The authors declare no competing financial interests.

