## Supplemental Materials Summary for "Cerebral Encoding of Word Classes is Distributed and Context-Dependent"

#### A. Supplementary methods

##### 1. Linguistic predictors

The distributions of the linguistic predictors word length, lexical frequency (9+$\log_{10}$(frequency)), surprisal (log10-transformed), entropy, and index across the three PoS categories (nouns, verbs and adjectives) are demonstrated in Figure 1. All metrics pertain to the content words that were actually presented to the participants during the recording sessions. The figure demonstrates that the surprisal and entropy values were higher in the scrambled condition, particularly for the nouns.

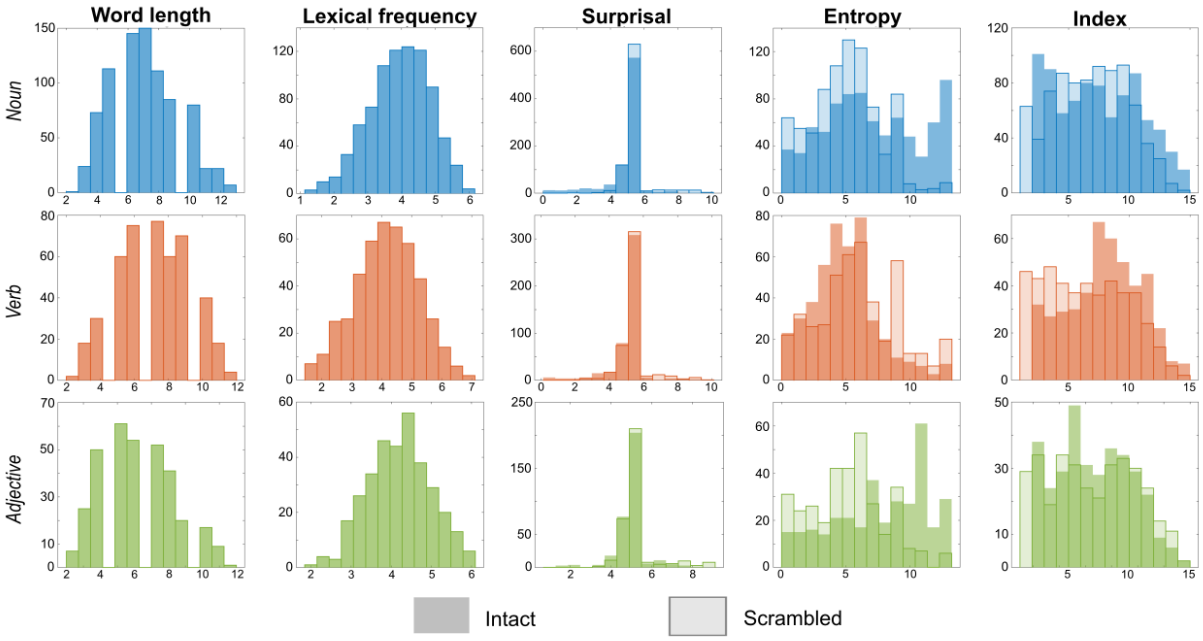


*Figure 1*. **Distribution of predictor variables by PoS category across conditions**: Distribution of word length, lexical frequency (9+log10(frequency), surprisal (log10-transformed), entropy, and index by Noun (blue), Verb (salmon), and Adjective (green) in the intact and scrambled conditions. The intact condition is shown in opaque colors and the scrambled condition in transparent colors with a contour.

We used multinomial logistic regression (MATLAB's *fitmnr* function) to investigate how linguistic variables predict PoS categories (noun, verb, adjective) in sentences and word lists. The full model included the following predictors: PoS~wordlength+lexfreq+index+surprisal+entropy. Model performance was assessed using log-likelihood, Akaike Information Criterion (AIC), and Bayesian Information Criterion (BIC). Model fit was better for sentences than for word lists (Sentences: Log-Likelihood = -2359.31, AIC = 4742.63, BIC = 4812.41; Word lists: Log-Likelihood = -2465.5065, AIC = 4955.01, BIC = 5024.79). To test whether sentence-level context improves PoS classification above and beyond word-level linguistic predictors, we fit a multinomial logistic regression model to the pooled sentence and word-list data:

PoS ~wordlength+lexfreq+index+surprisal+entropy+context,

with context indicating the presence or absence of sentence structure. PoS classification was most successful in sentence context (overall accuracy = 53.01%, balanced accuracy = 46.46%), slightly weaker for isolated word lists (52.48%, 43.52%), and did not improve when both contexts were combined (51.82%, 43.32%). This suggests that linguistic predictors capture PoS information more consistently in sentences than across mixed contexts, indicating that contextual structure contributes to PoS discriminability. The poor classification accuracy demonstrated that the relationship between the lexical/theoretical predictors and the PoS category cannot be modeled using a linear regression.

To evaluate the ability of lexical variables to predict PoS categories in a non-linear manner, we trained a multi-class support vector machine (SVM) classifier using a radial basis function (RBF) kernel. We used MATLAB's fitcecoc function with a one-vs-all coding scheme to support multi-class classification. This error-correcting output code (ECOC) framework decomposes the multiclass problem into binary classification subtasks. To account for class imbalance (44% percent of the content words were nouns), we computed inverse-frequency class weights and assigned them to training instances. Five-fold stratified cross-validation was used to ensure balanced representation of classes in each fold. Classification performance was evaluated using mean accuracy across the five folds. The random seed was fixed for reproducibility. This approach improved classification, particularly for the sentence condition (47.17 %) relative to the word list condition (38.79 %). A paired t-test confirmed the difference was statistically significant ($p<.05$). Figure 2 demonstrates classification performance for PoS categories from linguistic variables: (A) confusion matrix for the sentence condition; (B) confusion matrix for the word list condition; (C) difference matrix, i.e. the subtraction of word list from sentence (A–B), highlighting where classification accuracy improved (positive values) or worsened (negative values) in the sentence condition relative to the word lists. Color intensity (A, B) reflects classification percentages. The color scale in (C) encodes classification accuracy gain (yellow) vs classification accuracy loss (dark purple). Each matrix shows the percentage of predicted PoS labels (columns) given the true labels (rows) for three categories: Noun, Verb, and Adjective. Higher diagonal values indicate better classification accuracy. Off-diagonal values reflect misclassifications. Class-wise accuracy was highest for nouns (sentences: 51.6% vs word lists: 45.7%) and lowest for adjectives (sentences: 41.8% vs word lists: 23.4%). Adjectives showed the largest accuracy gain between conditions (18.4%), however, they also demonstrated the highest misclassification rates across PoS categories: in the sentence condition, out of 100 % of classifications, adjectives were misclassified as nouns in 33.8% and as verbs in 24.3%; in the word list condition, out of 100% classifications, adjectives were misclassified as nouns in 45.7% and as verbs in 30.9%.

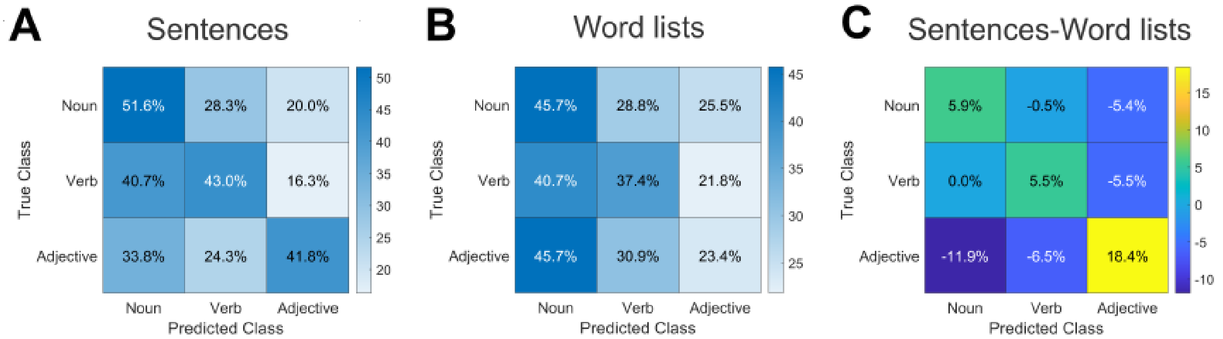


*Figure 2.* **Confusion Matrices:** (A) Confusion matrix for the intact sentence condition. (B) Confusion matrix for the scrambled condition. (C) Difference matrix of the type (A–B), highlighting classification accuracy gain (positive values, yellow color range) or loss (negative values, purple color range) in the intact condition relative to the scrambled one. Each matrix shows the percentage of predicted PoS labels (columns) given the true labels (rows) for three categories: Noun, Verb, and Adjective. Higher diagonal values indicate better classification accuracy. Off-diagonal values reflect misclassifications. Color intensity reflects classification percentages.

#### 2. Data acquisition and analysis

During the source reconstruction, source activity was projected onto 382 cortical parcels based on the Conte69 atlas. The following two files summarize (i) the atlas parcellation labels and (ii) the Brodmann areas (BA) and anatomical regions associated with the respective parcel indices:

(i) "Parcel_label.csv": the first column refers to the parcel index and the second column provides the parcellation label associated with this parcel index in the Conte69 atlas. The atlas parcellation labels are of the form: "L_6_B05_07", where L/R refers to the hemisphere; the number preceding B05 refers to the BA; the last number in the string refers to the ordinal position of the current parcels within the BA. Therefore, the parcel with label "L_6_B05_07" is the seventh parcel in the left BA 6.

(ii) "Brodmann_area_by_Parcel_index.pdf": The first column refers to the Brodmann areas, the second column provides the corresponding neuroanatomical region, the third column lists parcel indices for the left-hemispheric BA, while the fourth column lists parcel indices for the right-hemispheric BA. The rows of the table are color-coded by the cortical lobes: frontal (light-blue); temporal (magenta), parietal (green), occipital (orange), and midline (brown).

### B. Supplementary results

"Parcel_label_ordered_by_lobe_HM.csv": provides the mapping between the parcel indices used for visualization (raster plots) and the corresponding Conte69 atlas labels, together with hemisphere, Brodmann area, and anatomical lobe assignments.
