## Supplementary material for "Cerebral Encoding of Word Classes is Distributed and Context-Dependent": Brodmann Area by Parcel Index

| Brodmann_Are | neuroanatomical_region | Parc_index_LEFT | Parc_index_RIGHT |
| --- | --- | --- | --- |
| 1 | Middle primary somatosensory cortex | 40, 41, 42, 43 | 231, 232, 233, 234 |
| 2 | Posterior primary somatosensory cortex | 60, 61, 62, 63, 64, 65, 66 | 251, 252, 253, 254, 255, 256, 257 |
| 3 | Anterior primary somatosensory cortex | 37, 38, 39 | 228, 229, 230 |
| 4 | Primary motor cortex | 18, 19, 20, 21, 22, 23, 24, 25, 26, 27, 28, 29, 30, 31 | 209, 210, 211, 212, 213, 214, 215, 216, 217, 218, 219, 220, |
| 5 | Superior parietal lobule | 44, 45, 46 | 235, 236, 237 |
| 6 | Premotor cortex and Supplementary Motor Cortex (Secondary Motor Cortex) (Supplementary motor area) | 6, 7, 8, 9, 10, 11, 12, 13, 14, 15, 16, 17 | 197, 198, 199, 200, 201, 202, 203, 204, 205, 206, 207, 208 |
| 7 | Visuo-Motor coordination | 47, 48, 49, 50, 51, 52, 53, 54, 55, 56, 57, 58, 59 | 238, 239, 240, 241, 242, 243, 244, 245, 246, 247, 248, 249, 250 |
| 8 | (*includes* ) *Frontal eye fields* | 1, 2, 3, 4, 5 | 192, 193, 194, 195, 196 |
| 9 | Dorsolateral prefrontal cortex | 32, 33, 34, 35, 36 | 223, 224, 225, 226, 227 |
| 10 | Anterior prefrontal cortex (most rostral part of superior and middle frontal gyri) | 156, 157, 158, 159, 160 | 347, 348, 349, 350, 351 |
| 11 | Orbitofrontal area (orbital and rectus gyri, plus part of the rostral part of the superior frontal gyrus) | 162, 163, 164, 165, 166, 167, 168, 169 | 353, 354, 355, 356, 357, 358, 359, 360 |
| 12 | Orbitofrontal area (used to be part of BA11, refers to the area between the superior frontal gyrus and the inferior rostral sulcus) | | |
| 13 | Insular cortex | | |
| 16 | Insular cortex | | |
| 17 | Primary visual cortex (V1) | 173, 174, 175, 176, 177 | 364, 365, 366, 367, 368 |
| 18 | Secondary visual cortex (V2) | 178, 179, 180, 181, 182, 183, 184 | 369, 370, 371, 372, 373, 374, 375 |
| 19 | Associative visual cortex (V3, V4, V5) | 95, 96, 97, 98, 99, 100, 101, 102, 103, 104, 105, 106, 107, 108 | 286, 287, 288, 289, 290, 291, 292, 293, 294, 295, 296, 297, 298, 299 |
| 20 | Inferior temporal gyrus | 142, 143, 144, 145, 146, 147, 148 | 333, 334, 335, 336, 337, 338, 339 |

| | | | |
|---|---|---|---|
| 21 | Middle temporal gyrus | 127, 128, 129, 130 | 318, 319, 320, 321 |
| 22 | Part of the superior temporal gyrus, included in Wernicke's area | 114, 115, 116, 117, 118, 119, 120, 121, 122, 123 | 305, 306, 307, 308, 309, 310, 311, 312, 313, 314 |
| 23 | Ventral posterior cingulate cortex | 83, 84, 85 | 274, 275, 276 |
| 24 | Ventral anterior cingulate cortex | 152, 153, 154, 155 | 343, 344, 345, 346 |
| 25 | Ventromedial prefrontal cortex (subgenual area) | 161 | 352 |
| 26 | Ectosplenial portion of the retrosplenial region of the cerebral cortex | 190 | 381 |
| 27 | Presubiculum | 185 | 376 |
| 28 | Ventral entorhinal cortex | 188 | 379 |
| 29 | Retrosplenial cortex | 189 | 380 |
| 30 | Subdivision of retrosplenial cortex | 113 | 304 |
| 31 | Dorsal Posterior cingulate cortex | 67, 68, 69, 70 | 258, 259, 260, 261 |
| 32 | Dorsal anterior cingulate cortex | 149, 150, 151 | 340, 341, 342 |
| 33 | Part of anterior cingulate cortex | 191 | 382 |
| 34 | Dorsal entorhinal cortex (on the Parahippocampal gyrus) | | |
| 35 | Part of the perirhinal cortex (in the rhinal sulcus) | 187 | 378 |
| 36 | Part of the perirhinal cortex (in the rhinal sulcus) | 186 | 377 |
| 37 | Fusiform gyrus | 135, 136, 137, 138, 139, 140, 141 | 326, 327, 328, 329, 330, 331, 332 |
| 38 | Temporopolar area (most rostral part of the superior and middle temporal gyri) | 131, 132, 133, 134 | 322, 323, 324, 325 |
| 39 | Angular gyrus, considered by some to be part of Wernicke's area | 86, 87, 88, 89, 90, 91, 92 | 277, 278, 279, 280, 281, 282, 283 |
| 40 | Supramarginal gyrus considered by some to be part of Wernicke's area | 71, 72, 73, 74, 75, 76 | 262, 263, 264, 265, 266, 267 |
| 41 | Auditory cortex | 110, 111, 112 | 301, 302, 303 |
| 42 | Auditory cortex | 124, 125, 126 | 315, 316, 317 |
| 43 | Primary gustatory cortex | 93, 94 | 284, 285 |

| | | | |
|---|---|---|---|
| 44 | Broca's area, includes the opercular part and triangular part of the inferior frontal gyrus | 77, 78, 79, 80 | 268, 269, 270, 271 |
| 45 | Broca's area, includes the opercular part and triangular part of the inferior frontal gyrus | 81, 82 | 272, 273 |
| 46 | Dorsolateral prefrontal cortex | 170, 171, 172 | 361, 362, 363 |
| 47 | Orbital part of inferior frontal gyrus | 109 | 300 |
| 48 | Retrosubicular area (a small part of the medial surface of the temporal lobe) | | |
| 52 | Parainsular area (at the junction of the temporal lobe and the insula) | | |

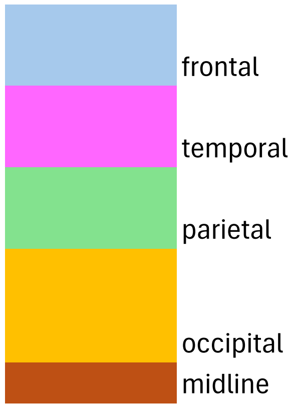
frontal
temporal
parietal
occipital
midline
